## Supplementary materials for "Robust integration of single-cell cytometry datasets"

### Supplementary Figures

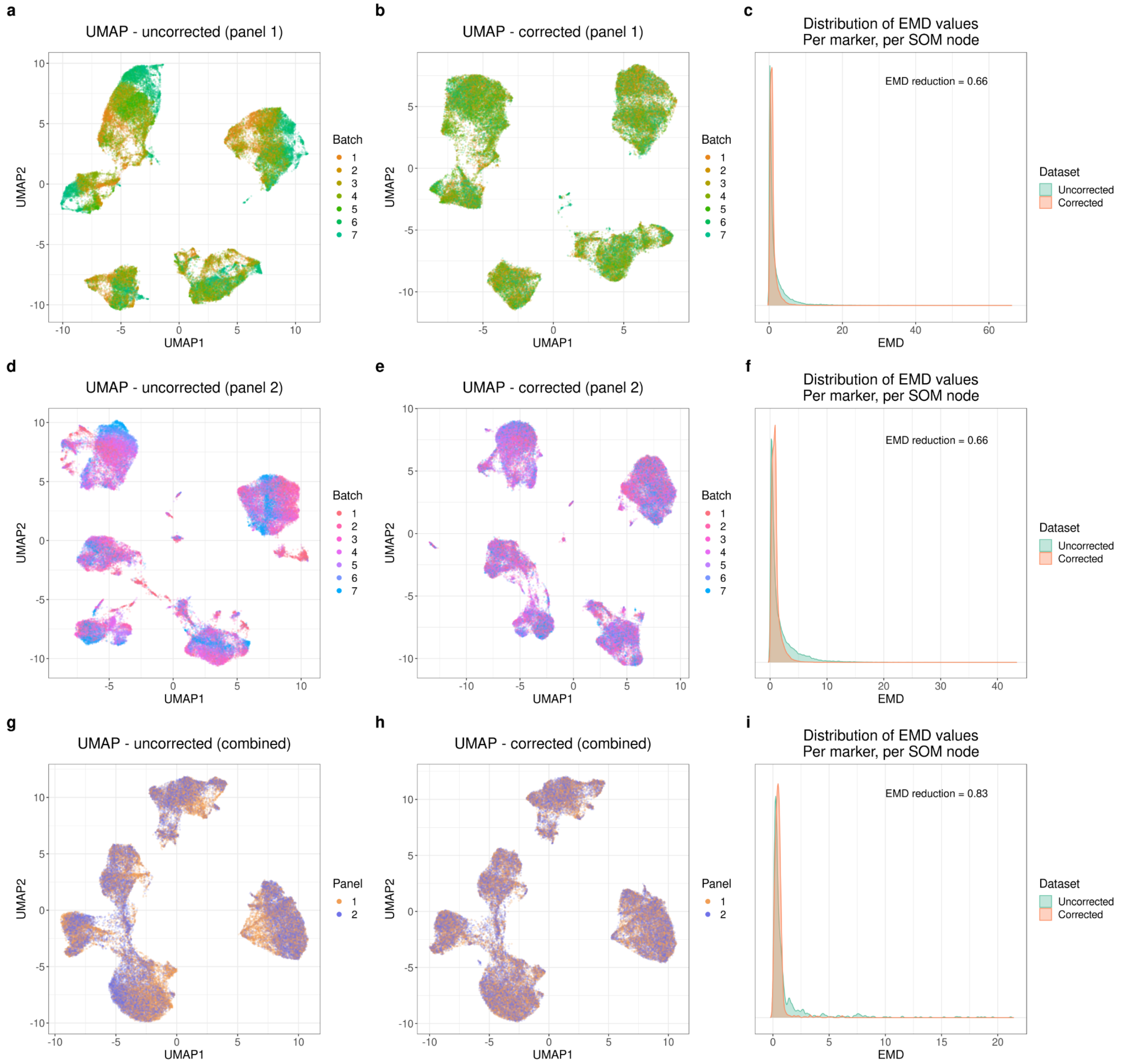


**Supplementary Figure 1.** Batch correction of the chronic lymphocytic leukemia (CLL) and healthy donor (HD) CyTOF dataset. **a-b**: UMAPs based on 36 markers for the panel 1 data before and after batch correction, generated using equal sampling of each batch to a total of 50,000 cells. **c**: Earth mover's distance (EMD) density plots for uncorrected and corrected data. The EMD reduction was 0.66 and the MAD score was 0.02. **d-f**: Same as **a-c** but for panel 2 and its 34 markers. The EMD reduction was 0.66 and the MAD score was 0.02. **g-i**: Same as **a-c** but for the co-batch correction of panels 1 and 2 and the 15 overlapping markers, using 25,000 cells per panel. The EMD reduction was 0.83 and the MAD score was 0.01.

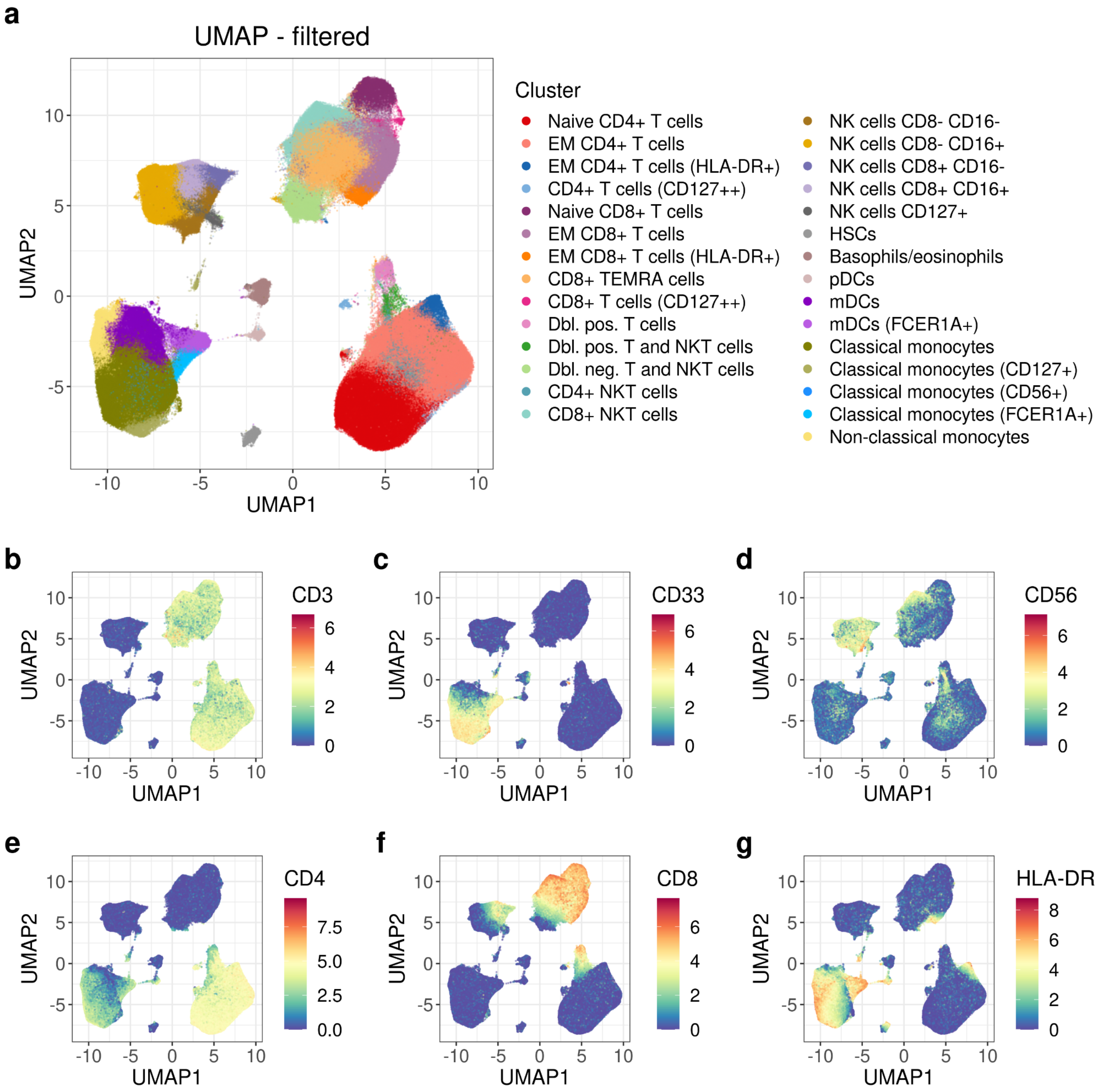


**Supplementary Figure 2.** Clustering results for the CLL and HD CyTOF dataset. **a**: UMAP for up to 2,000 cells from each of the 128 samples based on expression of the 23 clustering markers after removal of B, CLL, and poor-quality cells. **b-g**: Same as in **a**, but colored by expression of CD3, CD33, CD56, CD4, CD8, and HLA-DR.

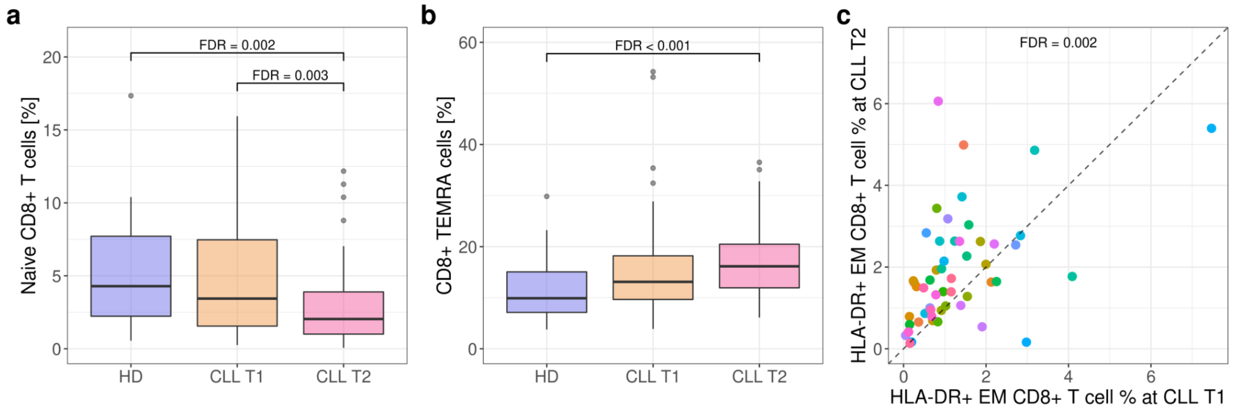


**Supplementary Figure 3.** Analysis of differential abundance within the T and NKT cell compartment in CLL and HD CyTOF data. **a-b**: Relative proportions of selected T and NKT cell populations for HD ($n$ = 20), CLL T1 ($n$ = 52), and CLL T2 ($n$ = 56) samples. FDR values provided for significant comparisons. **c**: Scatter plot for the relative proportions of the *paired* ($n$ = 52) CLL T1 and T2 patients in HLA-DR+ EM CD8+ T cells. FDR values provided for significant comparisons between CLL T2 vs. T1.

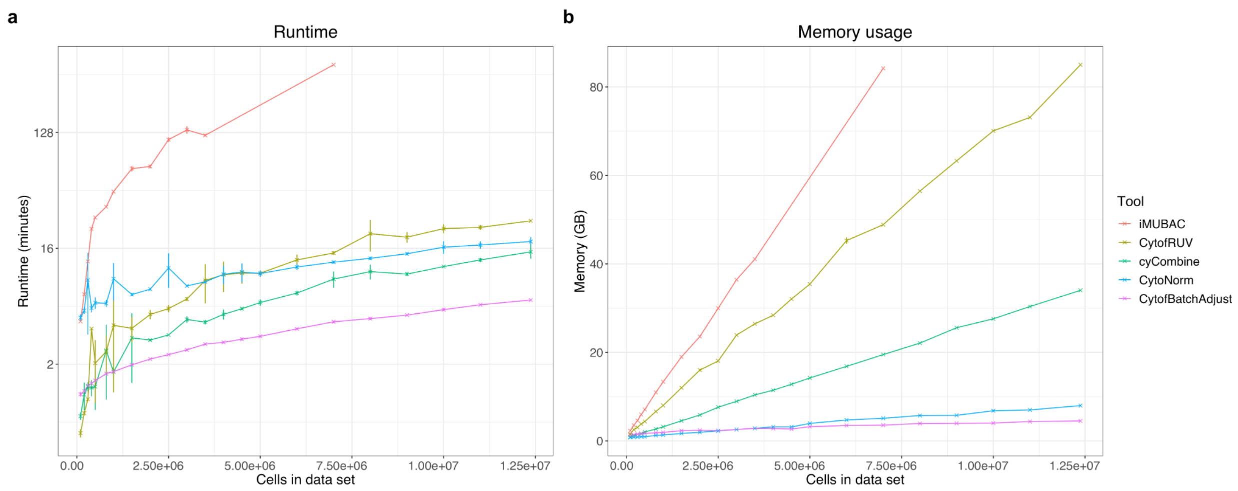


**Supplementary Figure 4.** Computational requirements of cyCombine and four other batch correction tools for a dataset with 38 markers and seven batches. **a**: Runtime in minutes (notice log scale on y axis) and **b**: Memory usage in GB. 40 cores and 100 gb memory was used for the system.

### Supplementary Discussion

**Panel merging**

Previous work by Abdelaal et al. (2019)[1] suggested approaches for 1) designing panels with an optimal overlap for imputation and 2) imputation based on k-Nearest Neighbors (kNN) and medians. CyTOFmerge[1] was developed for designing experiments that allow for integration of multi-panel data, but this has limited use for merging of pre-designed panels. Lee et al.[2] presented a flow cytometry method for merging purposes, but this works only for two files at a time, relies on domain knowledge, and has only been tested on lymphocyte data. Our work builds on these and demonstrates a more biologically intuitive approach for panel merging, which overcomes some of the limitations in the existing approaches. CytoBackBone[3] offers a solution more similar to the cyCombine panel merging module. For combining and integrating datasets, approaches like QFMatch[4], SIC[5], and MetaCyto[6] have also been presented. However, these are focused on combining the *results* of complete analyses and not on allowing truly integrated analysis from start to end. A thorough evaluation and discussion of the panel merging module of cyCombine and the existing alternatives is included in a vignette at https://biosurf.org/cyCombine.

**Analysis of CLL data**

Given that CLL can severely affect bone-marrow production of immune and hematopoietic cells[7,8], immune dysfunction in CLL is to be expected. The focus of our analysis was to identify features of the immune system that differentiate CLL patients from HDs and patients sampled at different times relative to treatment initiation. After integrating the two panels of the CLL dataset, we compared the overall frequency for each of the 29 populations in cells originating from panel 1 and panel 2, respectively. We observed a very strong Pearson correlation of 0.9996 making it reasonable to assume that any merged sample may be considered as a single combined sample for differential abundance analysis.

In this analysis, we found that, compared to HDs, close-to-treatment time point 2 (T2) CLL patients had higher amounts of T and NKT cells as well as HSCs (CD34+) (**Figure 2**). For HD vs. CLL time point 1 (T1), similar patterns were observed, but the difference in HSC frequencies was not statistically significant. However, when comparing CLL T2 vs. CLL T1 while accounting for the paired samples, the HSCs were significantly more abundant (logFC = 0.7) at CLL T2. Additionally, the most dramatic difference for HD vs. CLL T2 was also the HSCs (logFC = 1.2).

It is currently debated whether the absolute T cell counts in CLL are higher[9–11] or lower[12] than those found in HDs. Our results show a higher proportion of T and NKT cells in CLL patients, compared to HD, which is in line with the majority of the published works. Furthermore, the effects of CLL on the T cell compartment are also widely discussed[9–14], and in order to investigate the T and NKT cell compartment more deeply, we considered the populations as 'daughters' of their overall type, meaning that proportions were relative to the parent set of T and NKT clusters. This was done to account for the compositional nature of the data, which means that changes in overall cell type proportions can mask population-specific differences (when one population increases in frequency, the sum of frequencies of the rest of the populations will go down).

Within the group of T and NKT cells, we found that when comparing HD vs. CLL T1, the CLL samples had lower proportions of HLA-DR+ effector memory (EM) CD4+ T cells, and for CLL T2 vs. HD, we observed significantly lower levels of naive CD8+ T cells (**Supplementary Figure 3a**) and higher abundances of CD8+ terminally differentiated effector memory (TEMRA) cells in the CLL samples (**Supplementary Figure 3b**). Previous studies have also reported a general decrease in the naive T cell compartment for CLL patients[12,13]. Skews towards CD8+ EM T and TEMRA cells among CLL patients have also been reported[12,15], as well as a general increase in antigen-experienced[11] or memory T cells[13], as seen here, indicative of a low output of naive T cells.

Within the CLL patients, additional significant changes were detected. Overall, the two populations of HLA-DR+ EM CD8+ and CD4+ T cells constituted larger proportions of the total T and NKT cell compartment at T2 (**Figure 2f** and **Supplementary Figure 3c**), whereas the naive CD8+ T cells were less abundant at T2.

Elston et al. (2019)[13] associated a subpopulation of CD4+PD-1+HLA-DR+ T cells to progression in CLL, and specifically mentioned that this is most frequent in the EM compartment. PD-1+ expression patterns have also been discussed in relation to replicative senescence, which is associated with more aggressive disease in CLL patients[12]. In our cohort, we found PD-1 to be most highly expressed by the two HLA-DR+ EM T cell subsets, indicating that these clusters may actually encompass the PD-1+ fraction as well, supporting existing results. Within the CD4+ cluster, there was also a significantly higher median expression of PD-1 in CLL T2 samples compared to HDs (logFC = 0.47) (**Figure 2g**).

Taken together, our analysis of CLL patients shows that applying cyCombine to a multi-batch dataset enables co-analysis leading to the identification of characteristics commonly ascribed to the CLL immunophenotype, as well as novel cellular phenotypes only detectable when combining large panels.

**Benchmarking**

For data integration purposes, the total size of datasets may be very large. This means that that ideal batch correction tool has good scaling in terms of runtime and memory usage. We investigated this for the five tools we tested and compared to cyCombine (**Supplementary Figure 4**). All the tools scaled approximately linearly in their memory usage, with cyCombine showing medium usage. In terms of runtime, cyCombine scales approximately linearly and three of the other included tools have similar runtime scaling.
